# A Multisite Model of Allosterism for the Adenosine A1 Receptor

**DOI:** 10.1101/2020.10.14.338822

**Authors:** Giuseppe Deganutti, Kerry Barkan, Graham Ladds, Christopher A Reynolds

**Author notes:** New address: Sosei Heptares, Granta Park Steinmetz Building, Cambridge, CB21 6DG, UK.

## Abstract

Despite being a target for about one-third of approved drugs, G protein-coupled receptors (GPCRs) still represent a tremendous reservoir for therapeutic strategies against countless diseases. For example, several cardiovascular and central nervous systems conditions could benefit from clinical agents that activate the adenosine 1 receptor (A_1_R), however, the pursuit of A_1_R agonists for clinical use are usually impeded by both on- and off-target side effects. One of the possible strategies to overcome this issue is the development of positive allosteric modulators (PAMs) capable of selectively enhancing the effect of a specific receptor subtype and triggering functional selectivity (a phenomenon also referred to as bias). Intriguingly, besides enforcing the effect of agonists upon binding to an allosteric site, most of the A_1_R PAMs display intrinsic partial agonism and orthosteric competition with antagonists. To rationalize this behaviour, we simulated the binding of the prototypical PAMs PD81723 and VCP171, the antagonist 13B, and the bitopic agonist VCP746. We propose that a single PAM can bind several A_1_R sites rather than a unique allosteric pocket, reconciling the structure-activity relationship and the mutagenesis results.

## 1. Introduction

The four G protein-coupled receptors (GPCR) for adenosine (A_1_R, A_2A_R, A_2B_R and A_3_R) are involved in purinergic signalling^1^ in numerous tissues and organs^2^. Upon extracellular binding of the endogenous agonist to the orthosteric site, and intracellular recruitment of a preferential G protein, they produce a plethora of different effects through inhibition or stimulation of adenylyl cyclase. However, phospholipase Cβ, Ca^2+^, and the mitogen-activated protein kinases (MAPK) pathways are also relevant^3^.

The subtype A_1_R, which is widely expressed throughout the body, couples to the inhibitory G protein (G_i/o_) and is characterized by the highest affinity for adenosine (pKi ~ 7.0)^4^ amongst the ARs subtypes. In the central nervous system (CNS) it reduces the neuronal firing rate by blocking neurotransmitters release. It also mediates negative chronotropic and inotropic effects in the heart^5^; it inhibits lipolysis and the release of renin ^6^. The development of an A_1_R agonist could dramatically impact the treatment of central nervous system (CNS) and cardiovascular diseases^3,7^, however, both off-target (e.g. difficulties in achieving adequate A_1_R selectivity over the other ARs and other GPCRs)^8^ and on-target adverse effects (e.g. the undesired activation of a particular A_1_R signaling pathway in a given tissues or organ)^9,10^ have severely limited the clinical development of new agents.

Recently, the A_1_R structure in both inactive^11,12^ and active G_i2_-bound^13^ states has been determined. The A_1_R structural hallmark, like all other GPCRs, is a transmembrane domain (TMD) formed by seven α-helixes that span the cytosolic membrane from the extracellular N terminal to the intracellular C terminal and shapes the orthosteric and the intracellular G_i_ protein binding sites. Among the three A_1_R extracellular loops (ECLs), ECL2 is the longest and the most important for orthosteric ligand binding^14^. In both the active and inactive A_1_R conformations, the ECL2 helix orients almost perpendicularly to the plane of the membrane; this is distinct from the A_2A_R ECL2 helix, which is almost parallel to the membrane^11^. This topology divergence at the extracellular vestibule of ARs subtypes is believed to be responsible for the selectivity shown by positive allosteric modulators (PAMs) on A_1_R.

According to the functional definition, a PAM binds to a site other than the orthosteric one and increases the response triggered by the orthosteric agonist. However, this enhancing effect can be achieved in different ways^15,16^. On the muscarinic M_2_ receptor (M_2_R), for example, PAMs bind at the extracellular vestibule^17,18^, right over the orthosteric site, and increases the residence time (RT – the reciprocal of the dissociation constant, koff) of the agonists, thus slowing down their dissociation. Several PAMs of the glucagon-like peptide-1 receptor (GLP-1R)^19,20^, instead, make direct contact with the transmembrane helix 6 (TM6), stabilizing the receptor active conformation. Several A_1_R PAMs have been developed over recent years. Their action has been attributed to a secondary pocket on ECL2 that is able to cross-talk with the orthosteric site^21,22^. Targeting this allosteric site could favour the selective activation of A_1_R over the other AR subtypes, reducing the off-target adverse effects. Moreover, allosteric compounds can drive a functional selectivity, that is, activate a specific intracellular signalling path^23,24^, therefore reducing the A_1_R on-target adverse effects^25^.

Intriguingly, many 2-amino-3-benzoylthiophene A_1_R PAMs, like PD81723 and VCP171 (Figure 1), exert competitive orthosteric binding^26,27^ and intrinsic agonist activity^24,28^ at concentrations higher than required to enhance the orthosteric agonist affinities (which is usually 1-10 μM in vitro), indicating a possible propensity to bind to the orthosteric site, in addition to an allosteric site. The structure-activity relationship (SAR) of these compounds^29^ indicates the 2-amino group is essential for the allosteric activity, while carbonyl-containing substituents at the 3-position strongly support the effect, along with the benzoyl moiety and alkyl substituents in the 4- and 5-positions, possibly due to a secondary hydrophobic pocket within the allosteric site as proposed by Ijzerman and collaborators^29^. An intramolecular hydrogen bond between the 2-amino group and the adjacent carbonyl in the 3-position (suggested by the crystal structure of isolated 2-amino-3-benzoylthiophene analogues^30^) has been proposed as important for distinguishing between PAM and orthosteric competitive activity^30,31^. Indeed, the forced planarity introduced with the 2-aminothienopyridazines (i.e. compound 13B in Figure 1) generates antagonists or inverse agonists with activity in the low nanomolar range^30,32^. The unique pharmacological profile of the A_1_R PAMs stimulated the design of bitopic compounds (ligands able to simultaneously occupy both the orthosteric and the allosteric sites) to trigger biased agonism and potentially avoid on-target side effects. This is the case for VCP746 (Figure 1), which, besides an improved affinity relative to its individual orthosteric (adenosine moiety) and allosteric (VCP171) pharmacophores, showed biased agonism relative to prototypical A_1_R agonists^33^.

**Figure 1.**
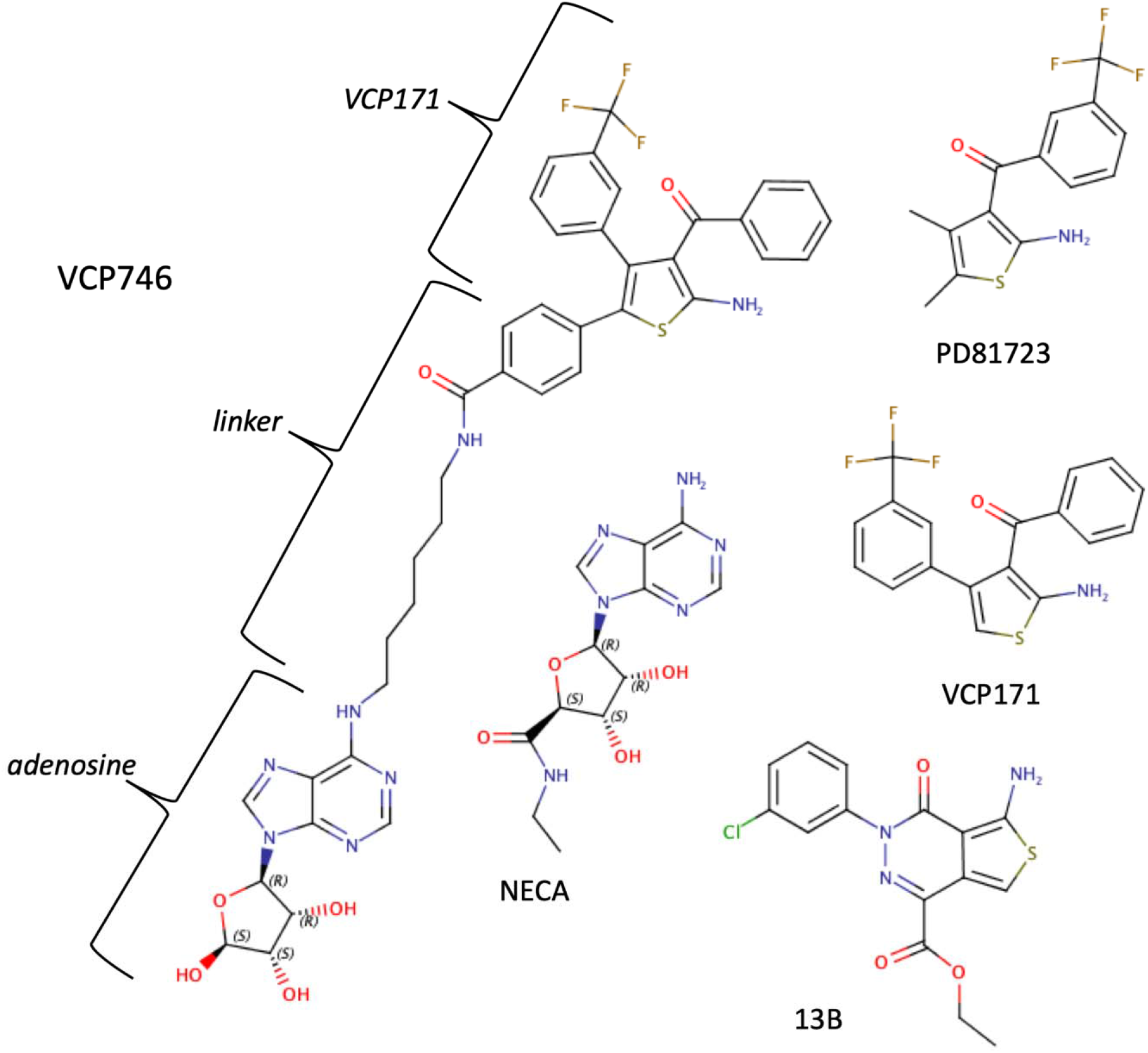
The A_1_R ligands considered in this study. PD81723 and VCP171 are positive allosteric modulators (PAMs) of the orthosteric agonist NECA. 13B is an A_1_R antagonist which is structurally related to the PAMs. VCP746 is a bitopic agonist formed from the endogenous agonist adenosine and the PAM VCP171, joined by an alkyl-aryl amide linker.

Even if extensive mutagenesis^21^ and computational^22^ work have profiled and characterised the A_1_R allosteric site, there is still uncertainty regarding the binding mechanism of both PAMs and bitopic A_1_R ligands and their SAR. In this study, we combined supervised molecular dynamics^34–37^ (SuMD) with more than 20 μs of classic MD to try rationalising part of the experimental body of data produced so far in literature. The simulated binding mechanism of the prototype PAMs PD81723 and VCP171 differed in presence or absence of the orthosteric agonist NECA, while the molecular planarity of compound 13B led to an antagonist-like binding profile. A possible kinetic model of allosterism is therefore proposed, along with the putative binding mode of the bitopic agonist VCP746 (Figure 1).

## 2 Methods

### 2.1 Experimental Methods

#### 2.1.1 Cell culture

CHO-K1-hA_1_R cells were maintained using standard subculturing routines as guided by the European Collection of Cell Culture (ECACC) and checked annually for mycoplasma infection using an EZ-PCR mycoplasma test kit from Biological Industries (Kibbutz Beit-Haemek, Israel). All procedures were performed in a sterile tissue culture hood using aseptic technique and solutions used in the propagation of each cell line were sterile and pre-warmed to 37°C. Cells were maintained at 37°C with 5% CO_2_, in a humidified atmosphere. This study used CHO cell lines as a model due to the lack of endogenous AR subtype expression (Brown *et al*., 2008). CHO-K1-A_1_R and CHO-K1 cells were routinely cultured in Hams F-12 nutrient mix (21765029, Thermo Fisher Scientific) supplemented with 10% Foetal bovine serum (FBS) (F9665, Sigma-Aldrich).

#### 2.1.2 cAMP accumulation assay

For cAMP inhibition experiments, CHO-K1-hA_1_R cells were harvested and re-suspended in stimulation buffer (PBS containing 0.1% BSA and 25 μM rolipram) and seeded at a density of 2,000 cells per well of a white 384-well Optiplate. CHO-K1-hA_1_R cells were then stimulated for 30 minutes with 1 μM forskolin, NECA and a range of PD81723 concentrations. cAMP levels were then determined using a LANCE^®^ cAMP kit as described previously^38^. To determine the A_1_R dependent effect of PD81723, CHO-K1-hA_1_R or CHO-K1 cells were stimulated with 1 μM forskolin and DMSO or 10 μM PD81723. Here, cAMP data is normalised to the maximum forskolin response 100 μM in each cell line. Data was globally fitted with the three-parameter logistical equation built into Graphpad Prism 8.0 and the operational model of allosterism and agonism^39^.

#### 2.1.3 Radioligand binding

100 μg protein per tube acquired from homogenisation of CHO-K1-hA_1_R cells in ice-cold buffer (2 mM MgCl_2_, 20 mM HEPES, pH 7.4) was used in radioligand displacement assays. The displacement of 1,3-[^3^H]-dipropyl-8-cyclopentylxanthine ([^3^H]-DPCPX), an A_1_R selective antagonist radioligand, at a fixed concentration (1 nM; around the Kd value (1.23 nM, as determined by saturation binding experiments) by increasing concentrations of NECA with DMSO control or PD81723 (10 μM). Non-specific binding was determined in the presence of 10 μM DPCPX. The binding affinity of NECA (pK_i_) was determined through fitting the ‘One site - Fit-Ki’ model. Statistical significance was determined compared to vehicle alone using Student un-paired T-test where * *p* < 0.05 was considered significant. Membrane and ligand was incubated for 60 minutes at room temperature in Sterilin™ scintillation vials (Thermo Fisher Scientific; Wilmington, Massachusetts, USA). Filtration through Whatman^®^ glass microfiber GF/B 25 mm filters (Sigma-Aldrich) separated free and bound radioligand and the radioactivity on each filter determined by the addition of 4 mL of Ultima Gold XR liquid scintillant (PerkinElmer), overnight incubation at room temperature and count using a Beckman Coulter LS 6500 Multi-purpose scintillation counter (Beckman Coulter Inc.; Indiana, USA).

### 2.2 Computational Methods

#### 2.2.1 Force field, ligands parameters and systems preparations

The CHARMM36^40,41^/CGenFF 3.0.1^42–44^ force field combination was employed in this work. Initial ligand (Figure 1) force field, topology and parameter files were obtained from the ParamChem webserver^42^. NECA is already well-parameterized in the CGenFF force filed. The dihedral terms of PD81723, VCP171, and VCP746 were visually inspected during short MD simulations in water and no further optimization was performed. The active state A_1_R coordinates were retrieved from the Protein Data Bank database^45^ entry 6D9H^13^. The A1 R intracellular loop 3 (ICL3) was modelled using Modeller 9.19^46^, the G_i_ protein α subunit helix H5 (residues 329 to 355) was retained in the intracellular cleft of the A_1_R in order to maintain the full-active conformation of the receptor, and the endogenous agonist adenosine was removed. Six systems were prepared for MD (Table 1): the PAM (PD81723 or VCP171) was placed about 40 Å away from the A_1_R in complex with NECA (inserted in the orthosteric site by superposition of PDB entry 2YDV^47^ onto 6D9H) in the extracellular bulk of the two different systems; one molecule (PD81723, VCP171, or VCP746) was placed about 40 Å away from the pseudo-apo A_1_R (obtained upon removing of the endogenous agonists) in the extracellular bulk of the three different systems. The inactive state A_1_R coordinates were retrieved from the Protein Data Bank database^45^ entry 5UEN^12^. After the fusion protein and the co-crystallized antagonist were removed, one molecule of compound 13B was placed about 40 Å away from the pseudo-apo A_1_R. For all six systems (Table 1), hydrogen atoms were added by means of the pdb2pqr^48^ and propka^49^ software (considering a simulated pH of 7.0); the protonation of titratable side chains was checked by visual inspection. The resulting receptors were separately inserted in a square 90 Å x 90 Å 1-palmitoyl-2-oleyl-sn-glycerol-3-phosphocholine (POPC) bilayer (previously built by using the VMD Membrane Builder plugin 1.1, Membrane Plugin, Version 1.1. at: http://www.ks.uiuc.edu/Research/vmd/plugins/membrane/), through an insertion method^50^, along with their co-crystallized ligand (and the water molecules within 5 Å of the ligand). The receptor orientation was obtained by superposing the coordinates on the corresponding structure retrieved from the OPM database^51^. Lipids overlapping the receptor transmembrane helical bundle were removed and TIP3P water molecules^52^ were added to the simulation box by means of the VMD Solvate plugin 1.5 (Solvate Plugin, Version 1.5. at <http://www.ks.uiuc.edu/Research/vmd/plugins/solvate/). Finally, overall charge neutrality was reached by adding Na^+^/Cl^−^ counter ions up to the final concentration of 0.150 M), using the VMD Autoionize plugin 1.3 (Autoionize Plugin, Version 1.3. at <http://www.ks.uiuc.edu/Research/vmd/plugins/autoionize/).

**Table 1.** Summary of all the simulations performed and the MD sampling time considered for the analysis. The binding of the PAMs PD81723 and VCP171 to the active state A_1_R was simulated both in the presence and absence of the orthosteric agonist NECA.

| **Ligand** | **Orthosteric NECA** | **# SuMD replica / path sampling** | **SuMD path sampling time** | **A1 conformational state** |
| --- | --- | --- | --- | --- |
| PD81723 | Yes | 10 | 2.34 μs | Active |
| PD81723 | No | 10 | 2.59 μs | Active |
| VCP171 | Yes | 10 | 2.40 μs | Active |
| VCP171 | No | 9 | 3.89 μs | Active |
| 13B | No | 11 | 6.50 μs | Inactive |
| VCP746 | No | 8 | 4.92 μs | Active |

#### 2.2.2 System equilibration and MD settings

The MD engine ACEMD^53^ was employed for both the equilibration and productive simulations. The equilibration was achieved in isothermal-isobaric conditions (NPT) using the Berendsen barostat^54^ (target pressure 1 atm) and the Langevin thermostat^55^ (target temperature 300 K) with low damping of 1 ps^-1^. A four-stage procedure was performed (integration time step of 2 fs): first, clashes between protein and lipid atoms were reduced through 2000 conjugate-gradient minimization steps, then a 2 ns long MD simulation was run with a positional constraint of 1 kcal mol^-1^ Å^-2^ on protein and lipid phosphorus atoms. During the second stage, 20 ns of MD simulation were performed constraining only the protein atoms, while in the last equilibration stage, positional constraints were applied only to the protein backbone alpha carbons, for a further 40 ns.

Productive trajectories (Table 1) were computed with an integration time step of 4 fs in the canonical ensemble (NVT). The target temperature was set at 300 K, using a thermostat damping of 0.1 ps^-1^; the M-SHAKE algorithm^56,57^ was employed to constrain the bond lengths involving hydrogen atoms. The cut-off distance for electrostatic interactions was set at 9 Å, with a switching function applied beyond 7.5 Å. Long-range Coulomb interactions were handled using the particle mesh Ewald summation method (PME)^58^ by setting the mesh spacing to 1.0 Å.

#### 2.2.3 SuMD and SuMD path sampling protocols

The supervised MD (SuMD) is an adaptive sampling method^59^ for speeding up the simulation of binding events between small molecules (or peptides^60,61^) and proteins^34,35^. Briefly, during the SuMD a series of short unbiased MD simulations are performed, and after each simulation the distances between the centres of mass (or the geometrical centres) of the ligand and the predicted binding site (collected at regular time intervals) are fitted to a linear function. If the resulting slope is negative (showing progress towards the target) the next simulation step starts from the last set of coordinates and velocities produced, otherwise, the simulation is restarted by randomly assigning the atomic velocities. To simulate the binding of PD81723, VCP171, and 13B to the A_1_R (Table 1, Videos S1-S5), the distance between the centroid of the small molecule and the centroid of the orthosteric residues N254^6.55^, F171^ECL2^, T277^7.42^, and H278^7.43^ was supervised during 500 ns long time windows until it reached a value less than 3 Å. In the case of VCP746 (Table 1, Video S6), the centroid of the adenosine portion of the bitopic agonist was considered. A further MD sampling protocol, namely SuMD path sampling (Table 1), was performed using the outputs from each SuMD replica. After alignment of the trajectory on the alpha carbon atoms of the protein, the MD frames characterized by protein-ligand contacts were divided into sequential groups according to the ligand RMSD to the starting positions (bin of 1 Å). Then, a frame from each of the groups (usually between 20 and 30) was randomly extracted and used as a starting point for classic MD simulations of 20 ns. SuMD path sampling allows wider exploration of macrostates identified by traditional SuMD, reducing the oversampling of microstates favoured by the supervision algorithm.

#### 2.2.4 MD Analysis

Only the output from the SuMD replica path sampling was considered for the analysis. Root mean square deviations (RMSD) were computed using VMD^62^. Interatomic contacts and ligand-protein hydrogen bonds were detected using the GetContacts scripts tool (https://getcontacts.github.io), setting a hydrogen bond donor-acceptor distance of 3.3 Å and an angle value of 150° as geometrical cut-offs. Contacts and hydrogen bond persistency are quantified as the percentage of frames (over all the frames obtained by merging the different replicas) in which protein residues formed contacts or hydrogen bonds with the ligand. Distances between atoms were computed using PLUMED 2.3^63^. The MMPBSA.py^64^ script, from the AmberTools17 suite (The Amber Molecular Dynamics Package, at http://ambermd.org/), was used to compute molecular mechanics energies combined with the generalized Born and surface area continuum solvation (MM/GBSA) method, after transforming the CHARMM psf topology files to an Amber prmtop format using ParmEd (documentation at <http://parmed.github.io/ParmEd/html/index.html). Water molecules with low mobility were detected using the AquaMMapS analysis^65^ on a 10 ns-long MD simulation of the apo A_1_R (coordinates were written every 50 ps of the simulation) restraining the backbone alpha carbon atoms similarly to the approach proposed in Wall *et al*^66^.

### 2.3 Numbering system

Throughout the manuscript, the Ballesteros-Weinstein residues numbering system for GPCRs^67^ is adopted.

## 3 Results and Discussion

### 3.1 PD81723 and VCP171 have a similar binding at the NECA-occupied A_1_R

The A_1_R-dependent effect of PD81723 (Figure 2a) was characterized by positive cooperativity on both NECA affinity (Figure 2b; NECA alone; pK_i_ - 6.74 ± 0.12; NECA + 10 μM PD81723; pKi - 7.194 ± 0.133; p < 0.05) and efficacy (Figure 2c; pK_B_ – 4.932 ± 0.21; log□β – 1.36 ± 0.31) without increasing the maximum response (Figure 2c). At a concentration of 10 μM, and in the absence of NECA, PD81723 triggered roughly 50% of A_1_R activation (Figure 2c). In line with the experimental conditions, we firstly simulated PD81723 and VC171 (Figure 1) in the presence of NECA bound to the orthosteric site of A_1_R (Video S1, Video S2). For both ligands, two ensembles of stable configurations (macrostates M1 and M2 in Figure 3a,b, 3e,f, Figure S2) were sampled in the cleft between the ECL2 helix and the β strand, while a further stable macrostate M3 (Figure 3a,b, 2e,f, Figure S2) was located between ECL2 and ECL3, near the top of TM2 and TM7.

**Figure 2.**
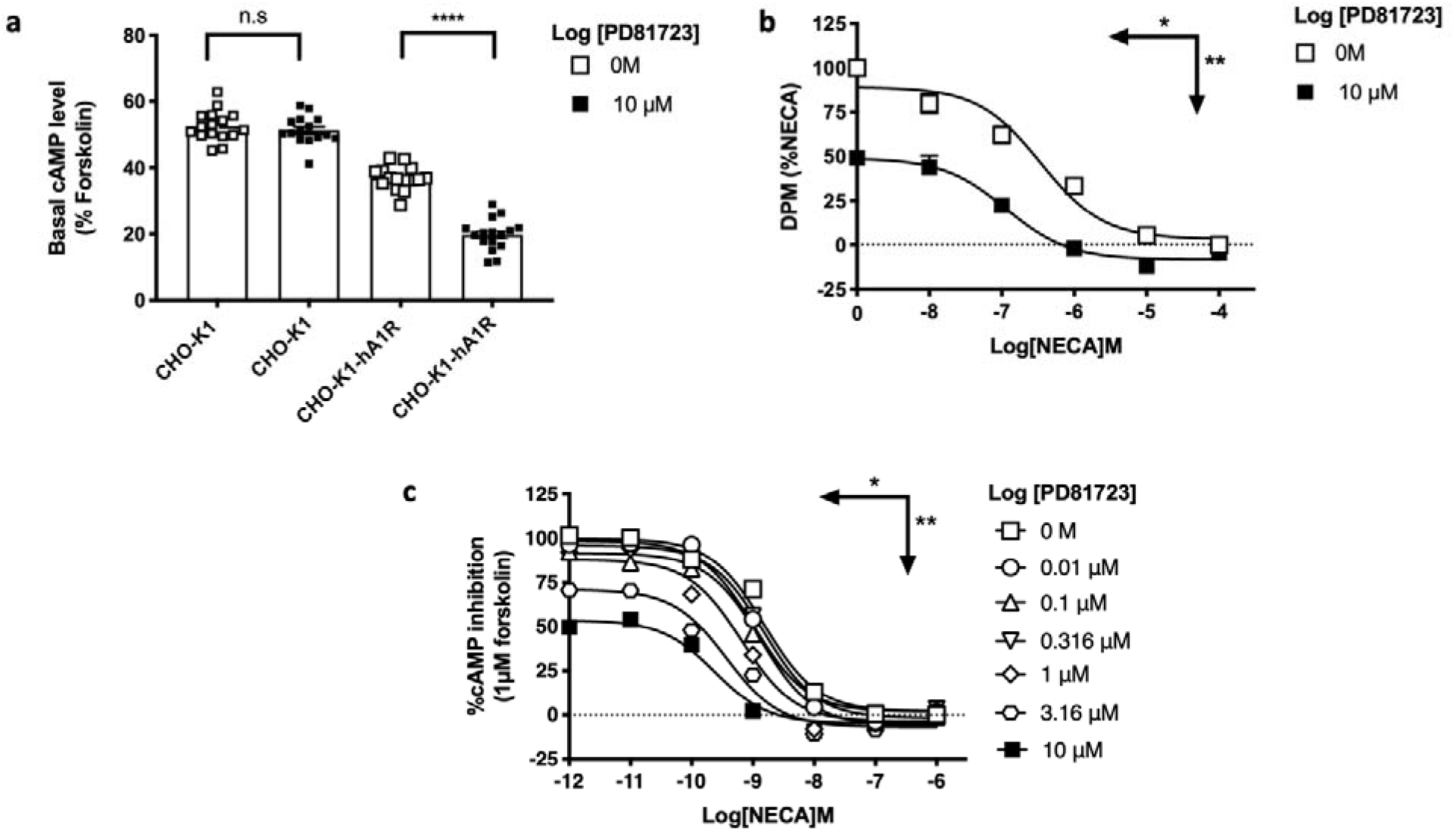
PD81723 exerts positive allosteric modulation on the NECA-bound A_1_R. **a)** A_1_R dependent effects of PD81723; CHO-K1 or CHO-K1-hA_1_R cells were stimulated with 1 μM forskolin and vehicle or vehicle containing 10 μM PD81723, data normalised to maximum forskolin response 100 μM; **b**) Affinity of NECA in the presence or absence of 10 μM PD81723; **c**) cAMP accumulation in the presence of increasing concentrations of PD81723. Data are mean ± SEM of at least 3 independent experiments, conducted in duplicate. Data was analysed using Student’s unpaired t-test where \**p* < 0.05; ** *p* < 0.01; *** *p* < 0.001; **** p < 0.0001 was considered significant.

**Figure 3.**
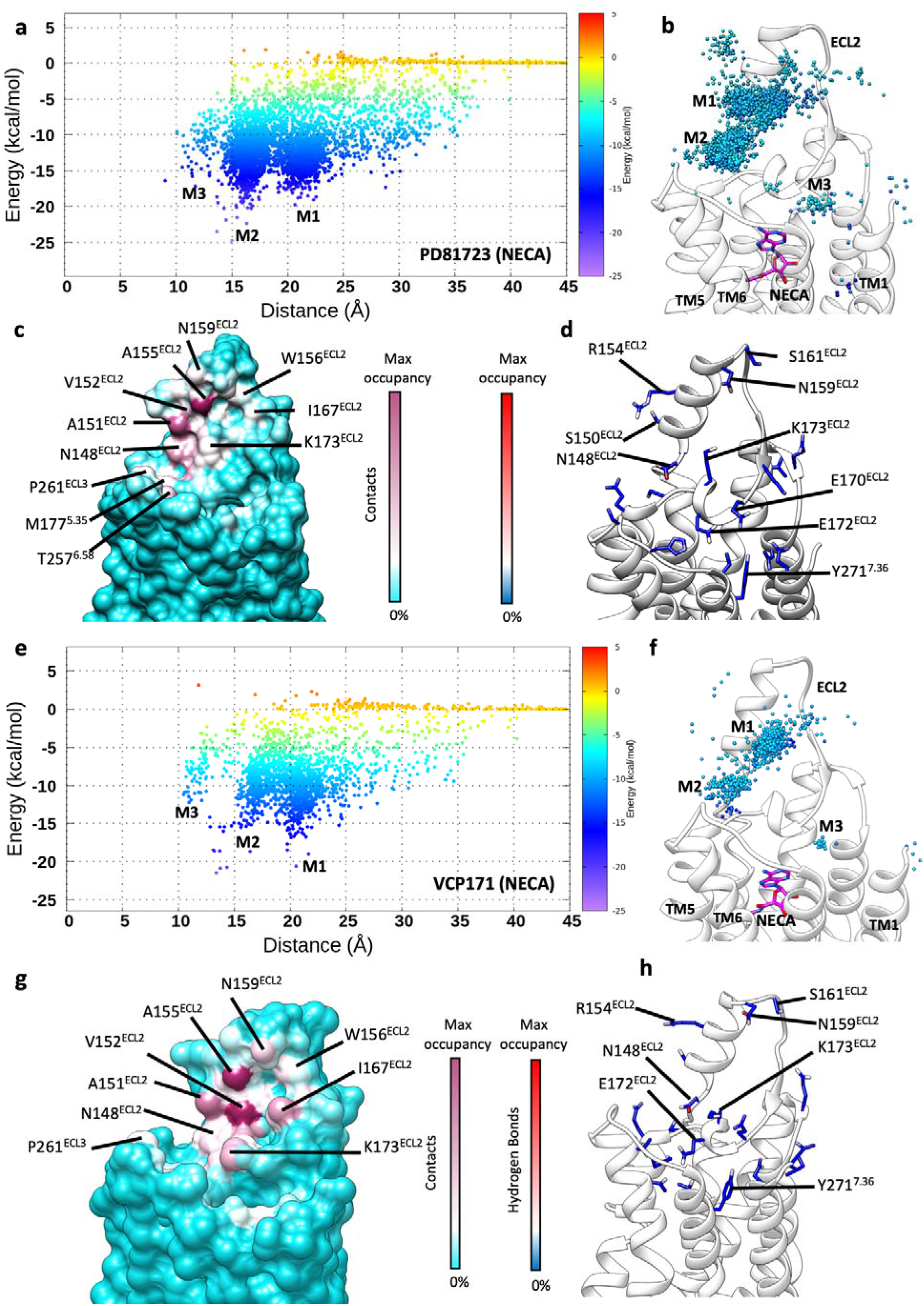
PD8172 and VCP171 SuMD binding in the occupied A_1_R (NECA bound to the orthosteric site). **a)** PD81723 binding energy landscape (the position of the metastable macrostates M1-M3 is reported); **b)** PD81723 centroid positions along the binding path, coloured according to the interactions experienced with A_1_R (energies higher than −5 kcal/mol are not shown); **c**) A_1_R - PD81723 contacts plotted on the receptor surface and coloured according to the occupancy (% MD frames in which a contact was present); **d**) A_1_R side chain atoms that formed hydrogen bonds with PD81723 and coloured according to the occupancy (% MD frames in which a contact was present; **e)** VCP171 binding energy landscape (the position of the metastable macrostates M1-M3 is reported)**; f)** VCP171 centroid positions along the association path, coloured according to the interactions experienced with A_1_R (energy higher than −5 kcal/mol are not shown); **g**) A_1_R – VCP171 contacts plotted on the receptor surface and coloured according to the occupancy (% MD frames in which a contact was present); **h**) A_1_R side chain atoms that formed hydrogen bonds with VCP171 and coloured according to the occupancy (% MD frames in which a contact was present).

The interaction analysis (Figure S1, Figure 3c,d, 2g,h) highlights the importance of residues A151^ECL2^, V152^ECL2^, A155^ECL2^, R154^ECL2^, W156^ECL2^, N159^ECL2^, V166^ECL2^, and I167^ECL2^ for stabilising the ligand at M1, while N148^ECL2^, E172 ^ECL2^, M177^5.35^, T257^6.58^, L258^6.58^ and P261^ECL3^ participated in the interaction network at M2. Both PD81723 and VCP171 established ECL2 a similar pattern of hydrogen bonds with A_1_R (Figure 3d,h), with the side chain of N148^ECL2^ highly engaged. VCP171 on the other hand, showed slightly more propensity to hydrogen bonding with N159^ECL2^ (Figure 3d, Figure 2h). Of note, E172^ECD^ bridges M2 to OS, interacting with both PD81723, VCP171 and NECA (Figure S2b,e). According to Nguyen *et al*^21^. N148^ECL2^A and K173^ECL2^A mutations decrease PD81723 binding cooperativity with NECA, while W156^ECL2^A causes a decrease in the cooperativity between VCP171 and NECA. Moreover, N148^ECL2A^A and I167^ECL2^A decrease the efficacy of both PD81723 and VCP171.

In the most stable microstate from M1 (which resembles the allosteric site recently proposed by Miao *et al*.^22^), residues A151^ECL2^, A155^ECL2^, W156^ECL2^, I167^ECL2^ and the alkyl chain of K173^ECL2^ formed hydrophobic contacts with the ligands (Figures S2a), while hydrogen bonds with S161^ECL2^, E172 ^ECL2^, N148^ECL2^ (PD81723 – Figure S2a-c), or K265^ECL3^ (VCP171– Figure S2d, Figure S3f) contributed to stabilising the PAMs in M2. Hydrophobic interactions with T270^7.35^, Y271^7.36^, I274^7.39^, the alkyl chain of K265^ECL3^, and a hydrogen bond with E170^ECL2^ instead stabilised the PAMs in M3 (the E170^ECL2^A mutation affects PD81723 binding cooperativity with NECA^21^; E170 borders the M3 site). Interestingly, macrostate M3 overlaps with the putative NECA unbinding path from the A2AR, as proposed by previous SuMD simulations^68^. The presence of a PAM right over the orthosteric site could be responsible for slowing down NECA dissociation^31,69^, as also determined for the class A M2R:iperoxo:LY211962 ternary complex^17^.

The ability of many A_1_R PAMs to reduce the orthosteric binding of antagonists^27,28,31,70^ could be due to the binding to distinct allosteric sites in the receptor inactive or active state^12^, The putative presence of more than one binding sites on the A_1_R vestibule (M1 to M3) may corroborate this hypothesis. M3, from this standpoint, resembles the secondary pocket observed in the inactive A_1_R structure^12^.

### 3.2 PD81723 and VCP171 bind to the unoccupied A_1_R with different mechanisms

The competitive binding, and the intrinsic activity exerted by several A_1_R PAMs, led us to hypothesise that they could also bind to the unoccupied orthosteric site, acting as partial agonists. We, therefore, simulated the binding of PD81723 and VCP171 in the absence of NECA (Video S3, Video S4). In contrast with the result obtained in the presence of NECA, the two ligands engaged the receptor with different patterns of interactions: VCP171 formed more contacts with TM1, TM3, TM3, and TM7 than PD81723, which instead mainly engaged ECL2 (Figure 4, Figure S1). In line with SuMD simulations in the presence of NECA (Figure 3), PD81723 formed transient metastable interactions with ECL2 (macrostates M1 and M2 in Figure 4a,b) and at the top of TM5 and TM6 (macrostate M3 in Figure 4a,b), before reaching the orthosteric site (macrostate OS in Figure 4a,b), The binding energy landscape (Figure 4a) shows that the most stable interactions with the protein were formed in M3, with OS characterised by slightly lower stabilisation. VCP171, on the other hand, was less prone to engage ECL2 in the intermediate macrostates M1 and M2 (Figure 4e,f), but instead populated the metastable M4 (Figure 4e,f), located between ECL2 and ECL3, near the top of TM1, TM2 and TM7, and the orthosteric site (Figure 4e,f).

**Figure 4.**
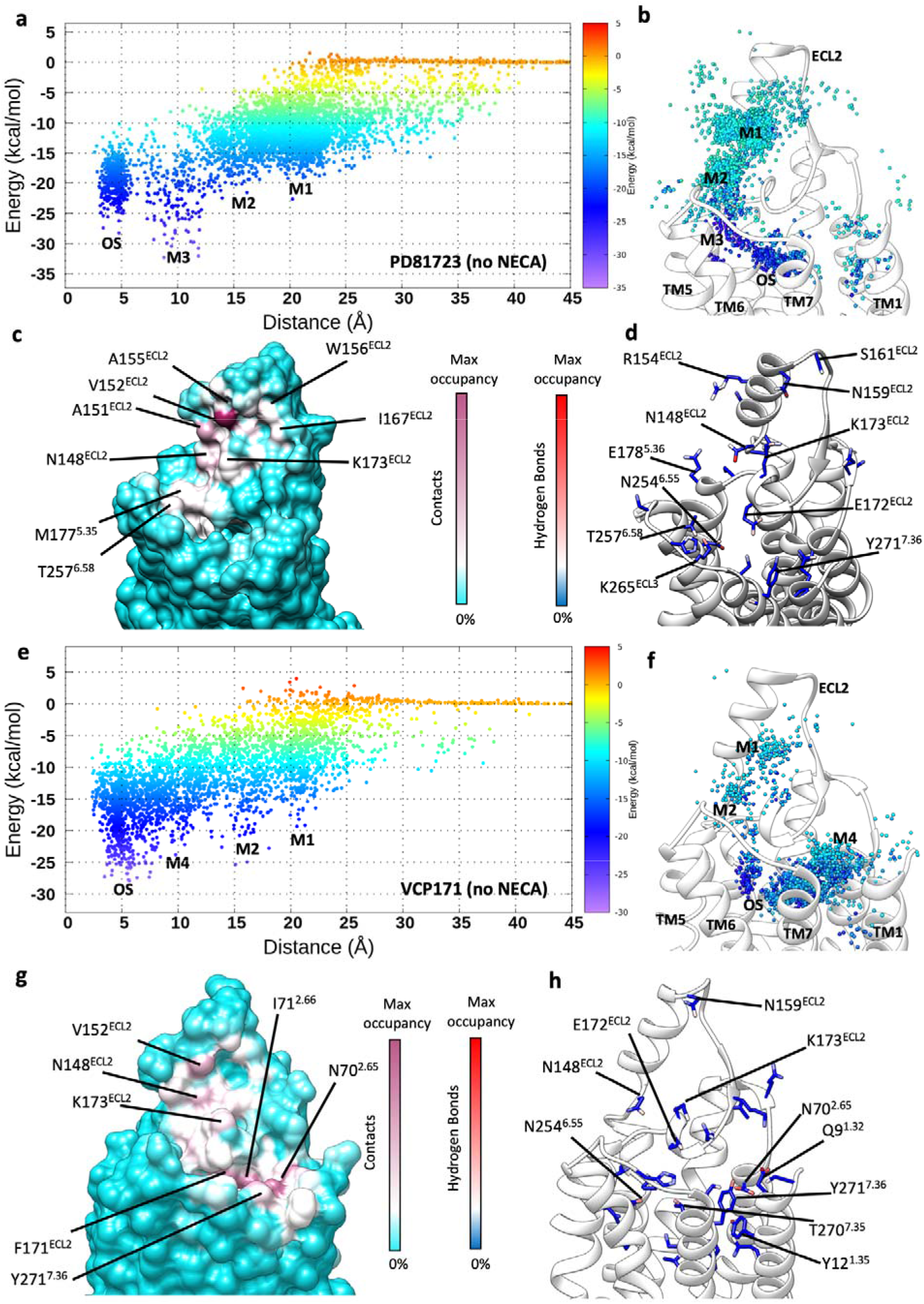
PD8172 and VCP171 SuMD binding in the unoccupied A_1_R. **a)** PD81723 binding energy landscape (the position of the metastable macrostates M1-M3 and the bound state OS is reported); **b)** PD81723 centroid positions along the binding path, coloured according to the interactions experienced with A_1_R (energies higher than −5 kcal/mol are not shown); **c**) A_1_R - PD81723 contacts plotted on the receptor surface and colored according to the occupancy (% MD frames in which a contact was present); **d**) A_1_R side chain atoms that formed hydrogen bonds with PD81723 and coloured according to the occupancy (% MD frames in which a contact was present; **e)** VCP171 binding energy landscape (the position of the metastable macrostates M1-M3 and the bound state OS is reported); **f)** VCP171 centroid positions along the association path, coloured according to the interactions experienced with A_1_R (energy higher than −5 kcal/mol are not shown); **g**) A_1_R – VCP171 contacts plotted on the receptor surface and coloured according to the occupancy (% MD frames in which a contact was present); **h**) A_1_R side chain atoms that formed hydrogen bonds with VCP171 and coloured according to the occupancy (% MD frames in which a contact was present).

In analogy with the occupied A_1_R, PD81723 formed hydrophobic interactions with A151^ECL2^, V152 ^ECL2^, A155^ECL2^, W156^ECL2^, I167^ECL2^ M177^5.35^, K173^ECL2^, and T257^6.58^, while hydrogen bonds were established with N148^ECL2^, S161^ECL2^, E172^ECL2^ (Figure 4c,d, Figure S1, Figure S3a-c). In the metastable macrostate M3 (Figure S3c), not sampled in the presence of NECA due to the partial overlap with the occupied orthosteric site, PD81723 formed a hydrogen bond with N254^6.55^ and hydrophobic interactions with M177^5.35^, T257^6.58^, F171^ECL2^ and L250^6.51^ (Figure 4c,d, Figure S1, Figure S3c). In the orthosteric site, PD81723 experienced numerous orientations, the most stable being driven by a hydrogen bond with S277^7.42^ and hydrophobic contacts with F171^ECL2^, L250^6.51^ and H278^7.43^ (Figure 4c,d, Figure S3d). While metastable macrostates M1 and M2 were preparatory for the productive orthosteric binding of PD81723 (through the macrostate M3 – Figure 4b), in the case of VCP171 these preliminary interactions appeared less important (Figure 4e), as it was much more prone to form stable intermediates M4 just over the orthosteric site prior to form the bound macrostate OS (Figure 4e,f). The A_1_R residues mostly involved in M4 were F8^1.31^, Q9^1.32^, Y12^1.35^, I71^2.66^, N70^2.65^, T270^7.35^ and T271^7.36^ (Figure 4g,h, Figure S1, Figure S3g), while in the orthosteric site V87^3.32^, N254^6.55^, L250^6.51^, I274^7.39^, F171^ECL2^, T277^7.42^ and H278^7.43^ were engaged (Figure 4h, Figure S1, Figure S3h).

The different binding energy landscape of the two PAMs (Figure 4a,e) suggest a different propensity to bind the A_1_R orthosteric site. Indeed, PD81723 has a SuMD energy profile characterised by configurations in macrostate (M3) slightly more stable than the orthosteric ones (OS) (Figure 4a), which to our experience could be indicative of a weak binder at the ARs^71^, while the gradual stabilisation gained by VCP171 along the binding path (Figure 4e) is consistent with a better binder. The different binding mechanisms exerted by PD81723 and VCP171 to the unoccupied A_1_R could be backed by the experimental evidence that receptor mutants at ECL2, except for E172^ECL2^A, affect (increase) only the affinity of PD81723, without significantly influencing VCP171. In light of these SuMD results, we propose that both VCP171 and PD81723 bind the orthosteric site of the unoccupied A_1_R triggering the experimentally observed partial activation of the receptor. The equilibrium shift towards active-like conformations would be consistent with the observed competitive displacement of antagonists^26,27^. A comparison with bound adenosine from the cryo-EM complex highlights several key orthosteric interactions formed by PD81723 (Figure S3i) and VCP171 (Figure S3l) that are consistent with the ones formed by the endogenous agonist.

Docking performed on the inactive human A_1_R^12^ showed that VCP171 could bind the A_1_R orthosteric site, overlapping the classical antagonist site. It assumes a conformation in which the 2-amino group and the 3-position carbonyl lie co-planar and forms a bidentate hydrogen bond with N254^6.55^ side-chain. The X-ray structure of an A_1_R PAM analogue showed the planar nature of these molecules^30^ in the solid phase. However, supramolecular interactions may force the planarity to optimize the intermolecular interactions and therefore the crystal lattice formation. Indeed, these SuMD simulations, which take into account a fully flexible hydrated environment, did not sample either a co-planarity between intramolecular donor and acceptor, or a bidentate hydrogen bond with N254^6.55^.

### 3.3 The introduction of planarity leads to an antagonist-like binding profile

The exploration of the SAR of the A_1_R PAMs led to the discovery of bicyclic and tricyclic derivatives acting as antagonists or inverse agonists, rather than positive allosteric modulators^30,32^. One of the most effective compounds, the 3-chlorophenylthienopyridazine compound 13B (Figure 1, Video S5), was active in the low nanomolar range^32^. To rationalise why the structural planarity introduced with the 9-membered ring changes the activity of these compound, we simulated the binding of compound 13B to the inactive A_1_R (Video S5, Figure 5).

**Figure 5.**
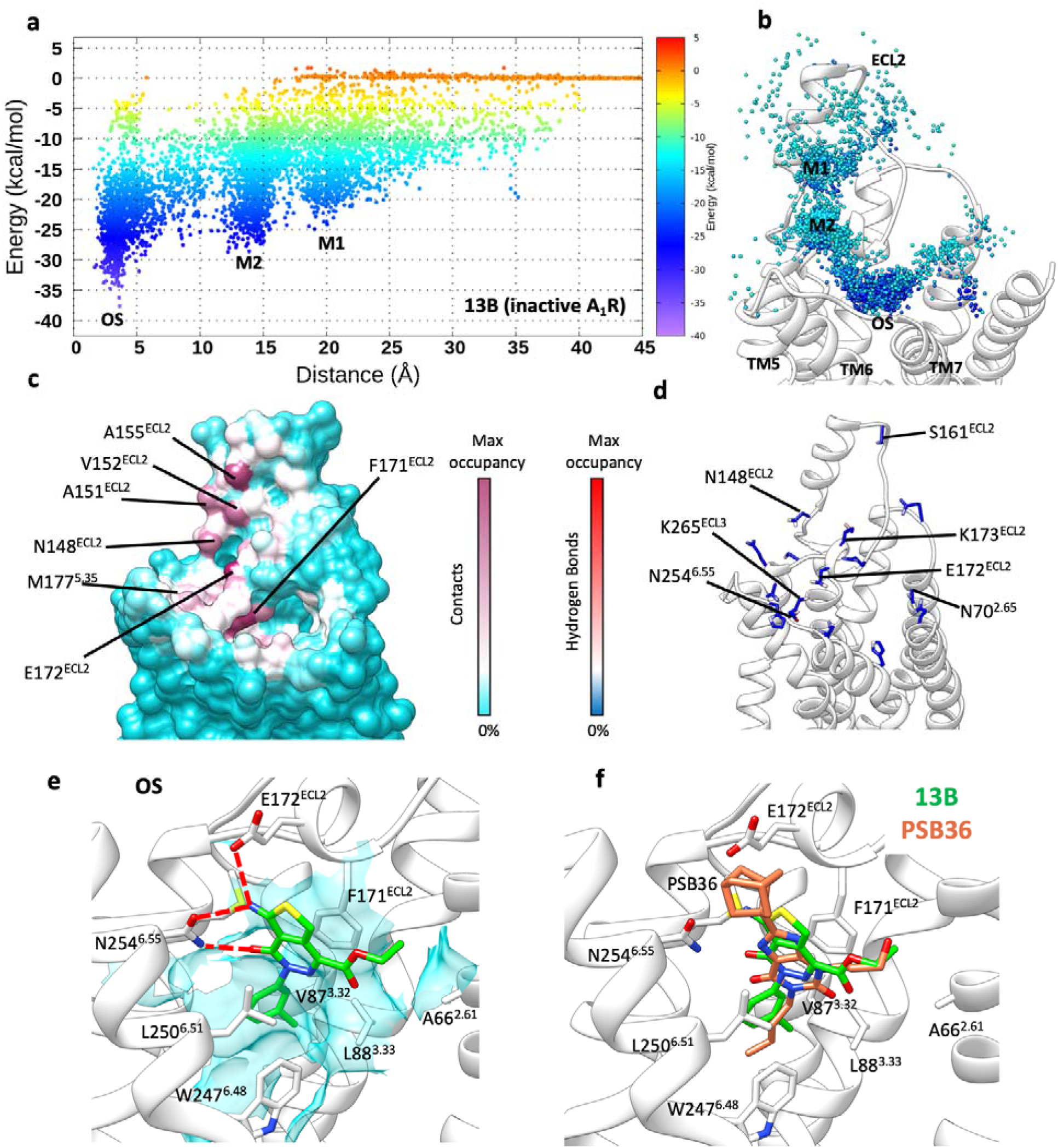
Compound 13B SuMD binding in the unoccupied inactive A_1_R. **a)** Compound 13B binding energy landscape (the position of the metastable macrostates M1-M2 and the bound state OS is reported); **b)** Compound 13B centroid positions along the binding path, coloured according to the interactions experienced with A_1_R (energies higher than −5 kcal/mol are not shown); **c**) A_1_R - compound 13B contacts plotted on the receptor surface and coloured according to the occupancy (% MD frames in which a contact was present); **d**) A_1_R side-chain atoms that formed hydrogen bonds with Compound 13B and coloured according to the occupancy (% MD frames in which a contact was present; **e)** representative bound conformation of compound 13B in the orthosteric states; **f)** binding mode comparison between compound 13B (green stick) and the selective antagonist PSB36 (tan stick). M3 is referred to in the text but there is no M3 in Fig. 5.

On the whole, by analogy with PD81723 and VC171, compound 13B’s interaction pattern (Figure S1) was characterised by two metastable macrostates on ECL2 (M1 and M2 in Figure 5a,b), and the orthosteric macrostate (OS in Figure 5a,b), which accounted for the higher stability. Intermolecular contacts with the protein were formed along the path with residues A151^ECL2^, V152^ECL2^, A155^ECL2^, E172 ^ECL2^, and M177^5.35^ (Figure 5c), while hydrogen bonds occurred with N148^ECL2^, K265^ECL2^, E172^ECL2^, and K265^ECL3^ side chains (Figure 5d). Representative configurations (microstates) from macrostates M1 and M2 (Figure S4) show the ligand laid on the surface of the loop, orienting the 3-chloropheny moiety towards the protein centre, and the ethyl ester towards the top of ECL2. Once 13B reached the orthosteric site, it experienced different orientations, hydrogen bonding with N254^6.55^ and E172^ECL2^ as well as engaging F171^ECL2^, V87^3.32^, L250^6.51^, W247^6.48^ in hydrophobic contacts (Figure S5ab). As expected, amongst the most stable OS configurations (Figure 5a), the inverse agonist engaged the receptor with the typical binding mode of the ARs antagonists, forming a bidentate hydrogen bond with N254^6.44^ (Figure 5e). The superposition with the A_1_R selective antagonists PSB36 (Figure 5f) shows the overlapping between compound 13B and the prototype antagonist, explaining the antagonist activity of the former and in line with earlier *in silico* docking experiments^12^.

So, why does compound 13B not have a modulatory effect on the active, agonist occupied A_1_R, since it is proposed to interact with the ECL2 region involved in the recognition of the PAMs PD81723 and VCP171? The reason does not reside in the interaction pattern with ECL2 side chains (which is very similar to the PAMs – Figures 2, Figure S1), but rather a possible explanation could lie in the interaction formed with the backbone of ECL2. Barely structured protein regions like the loops provide numerous hydrophilic spots for potential ligand interactions due to the solvent-exposed backbone oxygen and nitrogen atoms (Figure S6). While the 2-amino group (fundamental for PAM activity) of PD81723 and VCP171 hydrogen bonded with several oxygen atoms (Figure S7a-d), for compound 13B, the carbonyl oxygen atom in the 3-position mostly involved in a hydrogen bond with the M177^5.35^ backbone nitrogen atom. This difference is likely due to the different pharmacophore generated by the introduction of the planar 9-membered ring into the PAM active structure (Figure S8).

### 3.4 Exploring the bitopic ligand VCP476 binding mode

The pursuit of GPCR bitopic ligands is justified by the possibility of tuning the response triggered by the orthosteric ligand by combining both orthosteric agonist and (positive) allosteric pharmacophores in the same molecule. The resulting structure, indeed, should be able to concurrently engage the orthosteric and allosteric binding sites, thus stabilising a very specific conformation of the receptor^72,73^. The hetero-bivalent bitopic A_1_R ligand VCP746 (Figure 1, Video S6) is structurally formed by the endogenous agonist adenosine and the PAM VCP171. VCP746 showed bias towards the cAMP pathway relative to (probably G protein-dependent^74,75^) ERK1/2 signalling, and no observable effect on heart rate in isolated rat atria^33^. SAR studies^33,76^ determined that the 3-(trifluoromethyl)phenyl substituent is important for the allosterically-driven A_1_R bias, while the best linker length between the two pharmacophores (Figure 1) is six carbon atoms. Increasing the linker length to seven atoms produces bitopic ligands with increased bias profile and A_1_R selectivity over the other ARs, but a loss of affinity and potency.

Interestingly, the different SARs between bitopic ligands and PAMs suggests an alternative binding mode for the VCP746 allosteric pharmacophore relative to VCP171 alone. During SuMD binding simulations (Video S6, Figure 6), while the allosteric pharmacophore engaged ECL2 (Figure 6a,b), the adenosine portion of VCP171 was efficiently supervised to the orthosteric site, where it bound with the classic conformation and interaction pattern of AR agonists^47^ (Figure 6c,d,e). The bitopic ligand experienced the most stable configuration when the VCP746 adenosine part occupied the orthosteric site (macrostate OS in Figure 6a, Figure 5d, Figure 6e), suggesting the adenosine component of the ligand is particularly important for determining the affinity of the molecule. Focusing on the hydrogen bonds with A_1_R, VCP746 engaged numerous side chains (Figure S9) and the ECL2 backbone atoms (Figure S7f) as the other simulated ligands did. The simulations predicted that the linker would insert between the E172^ECL2^ and M177^5.35^ side chains and hydrogen bond with the ECL2 backbone (Figure 6e) thanks to the amide group bridging the aliphatic chain and the phenyl ring. During the preliminary steps of binding, the allosteric pharmacophore explored many orientations on ECL2 (Figure 6a,b), but eventually stabilised in a preferred binding mode near ECL2 as the adenosine reached the orthosteric site; the allosteric pharmacophore was partially constrained by the linker (BS in Figure 6a,b, Figure 6e,f). Overall, the VCP746 interaction pattern involved numerous interactions with E172^ECL2^ and K173^ECL2^ (Figure S1). These side chains, indeed, formed a sort of saddle for the allosteric pharmacophore, which often oriented the 3-(trifluoromethyl)phenyl group towards the hydrophobic pocket formed by the aliphatic chain of K173^ECL2^ and I167^ECL2^ (Figure 6f). This conformation diverges from the stable states sampled during SuMD binding of VCP171 to the occupied A_1_R, in which the PAM usually oriented the 3-(trifluoromethyl)phenyl moiety towards the top of ECL2 (Figure S2d). Some degree of similarity, on the other hand, is present between VCP171 metastable states during the binding to the unoccupied receptor (M1 in Figure S3e) and the conformation of the CP746 allosteric pharmacophore (Figure 6f).

**Figure 6.**
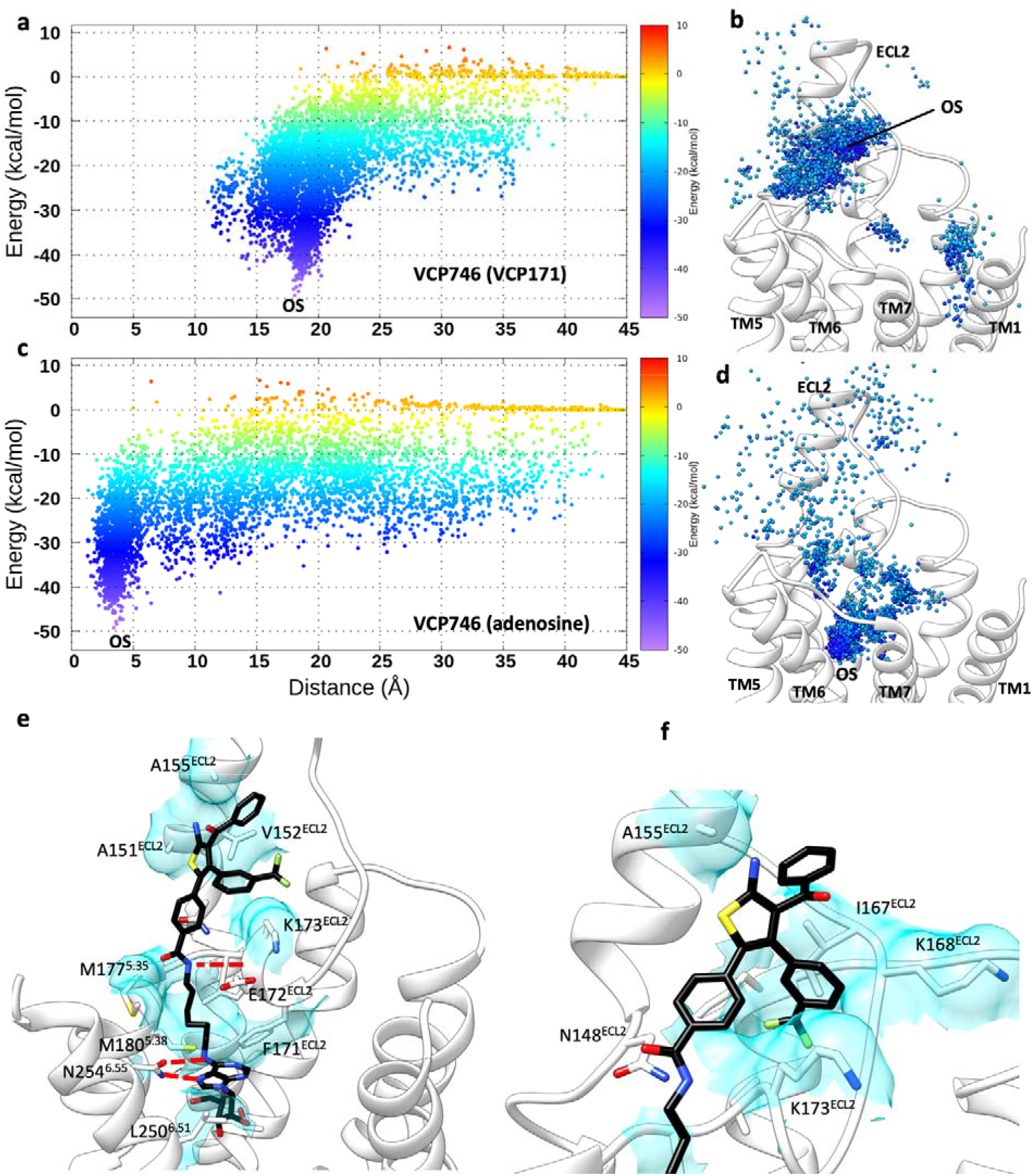
VCP746 SuMD binding in the unoccupied A_1_R. **a)** VCP746 binding energy landscape plotted on the distance between the allosteric pharmacophore (VCP171) and the orthosteric site (the most stable macrostate, OS, is indicated); **b)** centroid positions of the VCP746 allosteric pharmacophore (VCP171) along the binding path, coloured according to the interactions experienced with A_1_R (energies higher than −5 kcal/mol are not shown); **c**) VCP746 binding energy landscape plotted on the distance between the orthosteric pharmacophore (adenosine) and the orthosteric site (the most stable macrostates, OS, are indicated); **d)** centroid positions of the VCP746 orthosteric pharmacophore (adenosine) along the binding path, coloured according to the interactions experienced with A_1_R (energies higher than −5 kcal/mol are not shown); **e**) representative configuration from the stable macrostate OS showing the bitopic ligand engaged both the orthosteric site (hydrogen bond between adenosine and N254^6.55^) and ECL2; **f**) alternative stable conformation of the VCP746 allosteric pharmacophore on ECL2 resulting from rotation of the two substituted phenyl rings. VCP746 is shown as black stick, hydrogen bonds as dashed red lines and hydrophobic contacts as cyan transparent surfaces.

Taken together, these computational findings corroborate the hypothesis of a different binding mode for VCP171 as part of VCP746 and propose a sub pocket of ECL2 as involved in the observed bias^76^. Moreover, six carbon atoms could represent the best linker length for the affinity as it allows both a good orientation for the linker hydrogen bond with ECL2 and an adequate degree of conformational flexibility for the allosteric pharmacophore.

### 3.5 A multisite model reconciles divergent aspects of A1R PAMs pharmacodynamics

According to SuMD results, PD81723 and VCP171 interacted with the occupied A_1_R (e.g. NECA in the orthosteric site) in a similar way and formed stable ternary complexes with the two close and partially overlapping sites M1 and M2 on ECL2, and with M3 near the top of TM1, TM2 and TM7 (Figure 3a-d, Figure S2). M1 resembles the allosteric site proposed at the upper part of ECL2 by Yinglong *et al*.^22^, while M2 is located at the distal segment of ECL2, just over TM5 and the orthosteric site.

Interestingly, binding simulations to the unoccupied A_1_R showed both ligands able to form a final binary complex in the orthosteric site, consistent with their competitive behaviour and the intrinsic experimentally determined activity. However, in the unoccupied receptor, PD81723 conserved the same binding mechanism and formed roughly the same metastable states at ECL2 as it did in the occupied receptor (Figure 4a, Figure 3d, Figure S3), while VCP171 engaged A_1_R in a different way. Indeed, it poorly sampled the ECL2 metastable macrostates M1 and M2, preferring instead to engage the receptor at the M4 site, before entering into the orthosteric site (Figure 4a, Figure 3d, Figure S3). A different binding mechanism between PD81723 and VCP171 to the unoccupied A_1_R could explain mutagenesis experiment results showing that numerous ECL2 residues affect the affinity of PD81723, but not of VCP171^21^. A further intriguing aspect of the ECL2 mutants was the increased affinity measured for PD81723, except for the E172^ECL2^A A_1_R^21^ which decreased the pK_B_.

In light of all the considerations above, we propose a model with two sites (the orthosteric one and the putative, partially overlapping, allosteric sites M1 and M2 at ECL2) characterised by a similar (micromolar) affinity for the PAMs (Figure 7). As a general view, the binding at ECL2 would be kinetically favoured in consideration of the solvent exposure and the hydrophobic nature of the residues forming the pocket (that is, there should not be any hindrance or any stable water molecules network hampering the binding). Orthosteric complexes, on the other hand, would kinetically be less favoured due to the lower accessibility from the bulk. It follows that both PD81723 (Figure 7a,b) and VCP171 (Figure 7c,d) could bind the occupied A_1_R only engaging ECL2 (and M3 to a lesser extent) to exert PAM activity. On the unoccupied receptor, while PD81723 (Figure 7b) easily forms preliminary complexes with A_1_R ECL2 before engaging also the orthosteric site, VCP171 (Figure 7c) could form the orthosteric binary complex through M3 rather than via the ECL2 intermediate states M1 and M2. In this speculative scenario, mutations destabilising the PD81723:ECL2 complexes would paradoxically increase the measured affinity by modifying the binding pathway and shifting the equilibrium towards the orthosteric states (with the exception of E172^ECL2^, which is involved also in orthosteric interactions). This effect would be less significant for VCP171, which tends to engage the unoccupied A_1_R through a different path. Compound 13B showed a binding profile in line with the experimental behaviour. Indeed, it engaged the inactive A_1_R consistently with as a classic orthosteric antagonist, as it formed stable complexes with the same fingerprints of well-known ARs antagonist (Figure 5a,b). Also, compound 13B sampled kinetically accessible metastable states at ECL2 before reaching the final orthosteric complex (Figure 7e). However, its planarity affected the interaction pattern with the ECL2 sites, resulting in an unfavourable geometry for PAM activity.

**Figure 7.**
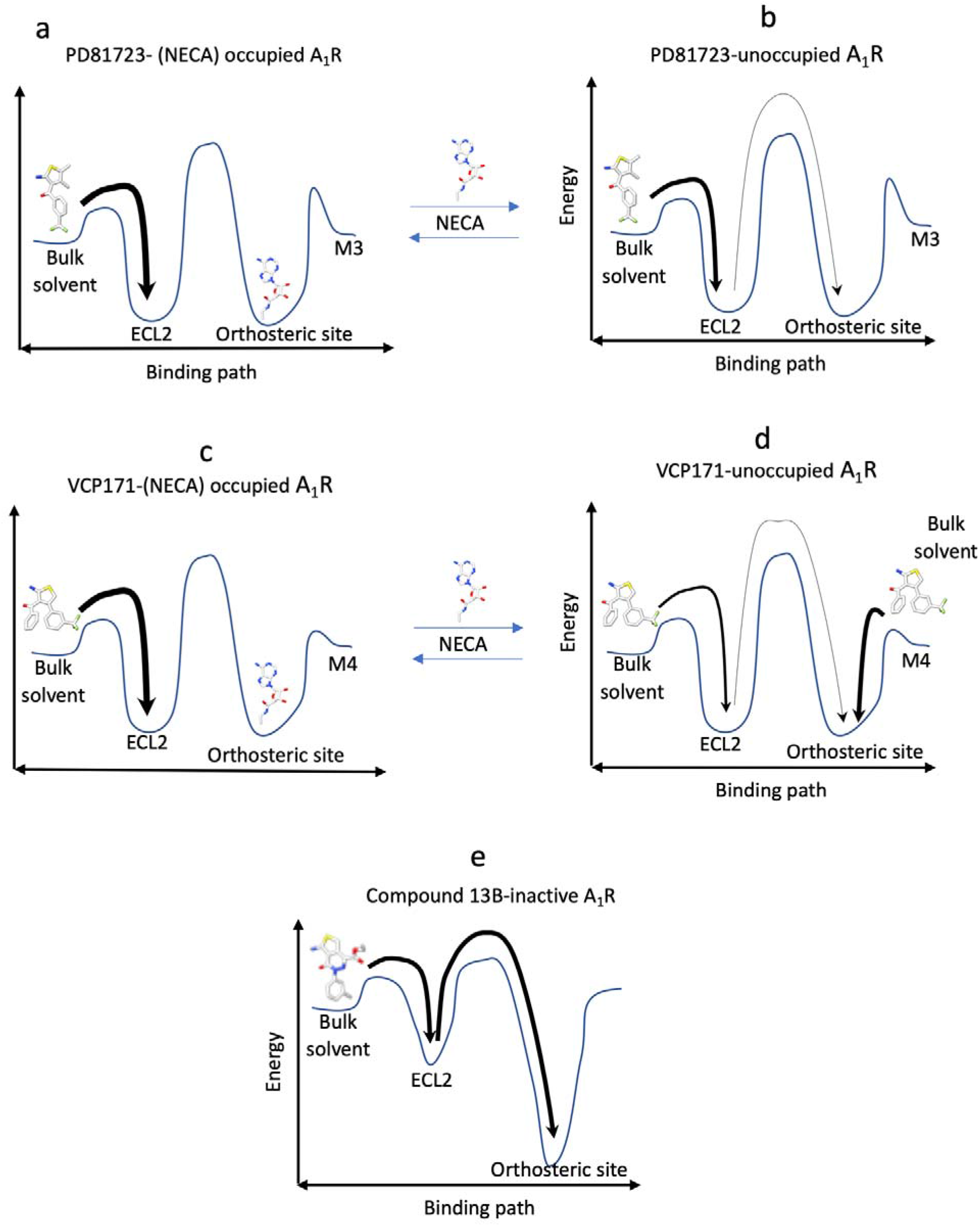
Scheme of the proposed binding model for A_1_R PAMs. **a)** On the occupied A_1_R, PD81723 binds to ECL2; **b)** on the unoccupied A_1_R, PD81723 binds to ECL2 first, then the orthosteric site; **c)** on the occupied A_1_R, CP171 binds to ECL2; **d)** on the unoccupied A_1_R, VCP171 can bind directly from the metastable state M3, and to a lesser extent from the ECL2 site; **e**) compound 13B has a binding mechanism that diverges from both of the two PAMs.

SuMD simulations have indicated how small structural modifications could have a large effect on binding mechanisms. Notably, we don’t univocally indicate a precise site triggering the allosteric effect on A_1_R (even if one of M1, M2 and M3/M4 could be the main site responsible for the effect) but rather we describe a more dynamic model in which more sites could contribute to engaging the PAMs. From this standpoint the bitopic ligand VCP746 fits well into this proposed functional plasticity, as the allosteric pharmacophore was proposed to bind with a geometry alternative to the one characterising VCP171 alone, but still capable of triggering a biased response upon orthosteric binding of the adenosine pharmacophore of the molecule.

This hypothesis could extend to other allosteric modulators that bind to the GPCR extracellular vestibule. Intriguingly, these sites could also be involved in the binding of orthosteric ligands, as has been experimentally proposed for the A_1_R^14^. An allosteric modulator whose binding site overlaps the binding or unbinding path of an orthosteric ligand could give rise to a complex scenario characterised by reciprocal competition for metastable sites, with consequences depending on the ligands kinetics and the balance between alternative (un)binding paths. This picture may partially explain a conflicting aspect of GPCR allostery^77^, namely why similar orthosteric ligands can respond very differently to the same PAM or NAM (a phenomenon called probe dependency). Indeed, in the simplest view, if an orthosteric ligand has to compete with a modulator for metastable sites along the binding or unbinding path (which in turn can be heavily affected by small structural changes in the ligand), then its affinity for the receptor could respectively decrease or increase in response to the modified (un)binding paths, with repercussion on the activity.

## 4 Conclusion

We proposed a multisite binding model for the prototypic PAMs of A_1_R. Instead of locating one distinct pocket on ECL2 that is putatively responsible for the allosteric effect, simulations suggested more extracellular sites able to bind the ligands in accordance with the SAR. In the absence of an agonist, the intrinsic agonism displayed by the PAMs could be due to some degree of orthosteric binding rather than a long-range effect triggered on the receptor conformation from an allosteric pocket(s). Interestingly, despite the structural similarity displayed by PD81723, VCP171, and 13B, divergent binding paths were observed to the apo A_1_R. This suggests a dramatic influence exerted by small chemical modifications on the binding kinetics.

Simulations of VCP746, to the best of our knowledge the first example of dynamic docking of a GPCR bitopic ligand, showed that linking a PAM to an agonist can modify the binding characteristic of the former, probably explaining the different pharmacology in terms of signalling and bias. The findings here reported help in understanding the binding mode of PAMs and bitopic ligands and therefore representing a further step towards the rational development of efficacious therapeutic targeting A_1_R.

## ASSOCIATED CONTENT

**Supporting Information (PDF)**

**Supplementary Videos S1-S6 (mp4)**.

## AUTHOR INFORMATION

### Present Addresses

§ New address: Sosei Heptares, Granta Park Steinmetz Building, Cambridge, CB21 6DG, UK

### Author Contributions

The manuscript was written through contributions of all authors. All authors have given approval to the final version of the manuscript.

## ACKNOWLEDGMENT

CAR is grateful for a Royal Society Industry Fellowship.

