## Supporting Information (PDF) for "A Multisite Model of Allosterism for the Adenosine A1 Receptor"

**TABLE of CONTENTS**

Supplementary Video S1-S6 captions pg. 1,2

Supplementary Figures S1-S9 pg. 3-11

**Supplementary Video S1.**

**SuMD simulations of PD81723 binding to the (NECA) occupied $A_1R$.** PD81723 and NECA are shown in van der Waals representation, while the protein residues in its proximity are shown as stick representation. The main hydrogen bonds are highlighted with red dotted lines. The simulation time indicated is cumulative of all the ten replicas. SuMD path sampling was seeded from these SuMD simulations.

**Supplementary Video S2.**

**SuMD simulations of VCP171 binding to the the (NECA) occupied $A_1R$.** VCP171 and NECA are shown in van der Waals representation, while the protein residues in its proximity are shown as stick representation. The main hydrogen bonds are highlighted with red dotted lines. The simulation time indicated is cumulative of all the ten replicas. SuMD path sampling was seeded from these SuMD simulations.

**Supplementary Video S3**

**SuMD simulations of PD81723 binding to the apo $A_1R$.** PD81723 is shown in van der Waals representation, while the protein residues in its proximity are shown as stick representation. The main hydrogen bonds are highlighted with red dotted lines. The simulation time indicated is cumulative of all the ten replicas. SuMD path sampling was seeded from these SuMD simulations.

**Supplementary Video S4**

**SuMD simulations of VCP171 binding to the apo $A_1R$.** VCP171 is shown in van der Waals representation, while the protein residues in its proximity are shown as stick representation. The main hydrogen bonds are highlighted with red dotted lines. The simulation time indicated is cumulative of all the ten replicas. SuMD path sampling was seeded from these SuMD simulations.

**Supplementary Video S5**

**SuMD simulations of 13B binding to the apo inactive $A_1R$.** 13B is shown in van der Waals representation, while the protein residues in its proximity are shown as stick representation. The main hydrogen bonds are highlighted with red dotted lines. The simulation time indicated is cumulative of all the ten replicas. SuMD path sampling was seeded from these SuMD simulations.

**Supplementary Video S6**

**SuMD simulations of VCP746 binding to the apo $A_1R$.** VCP746 is shown in van der Waals representation, while the protein residues in its proximity are shown as stick representation. The main hydrogen bonds are highlighted with red dotted lines. The simulation time indicated is cumulative of all the ten replicas. SuMD path sampling was seeded from these SuMD simulations.

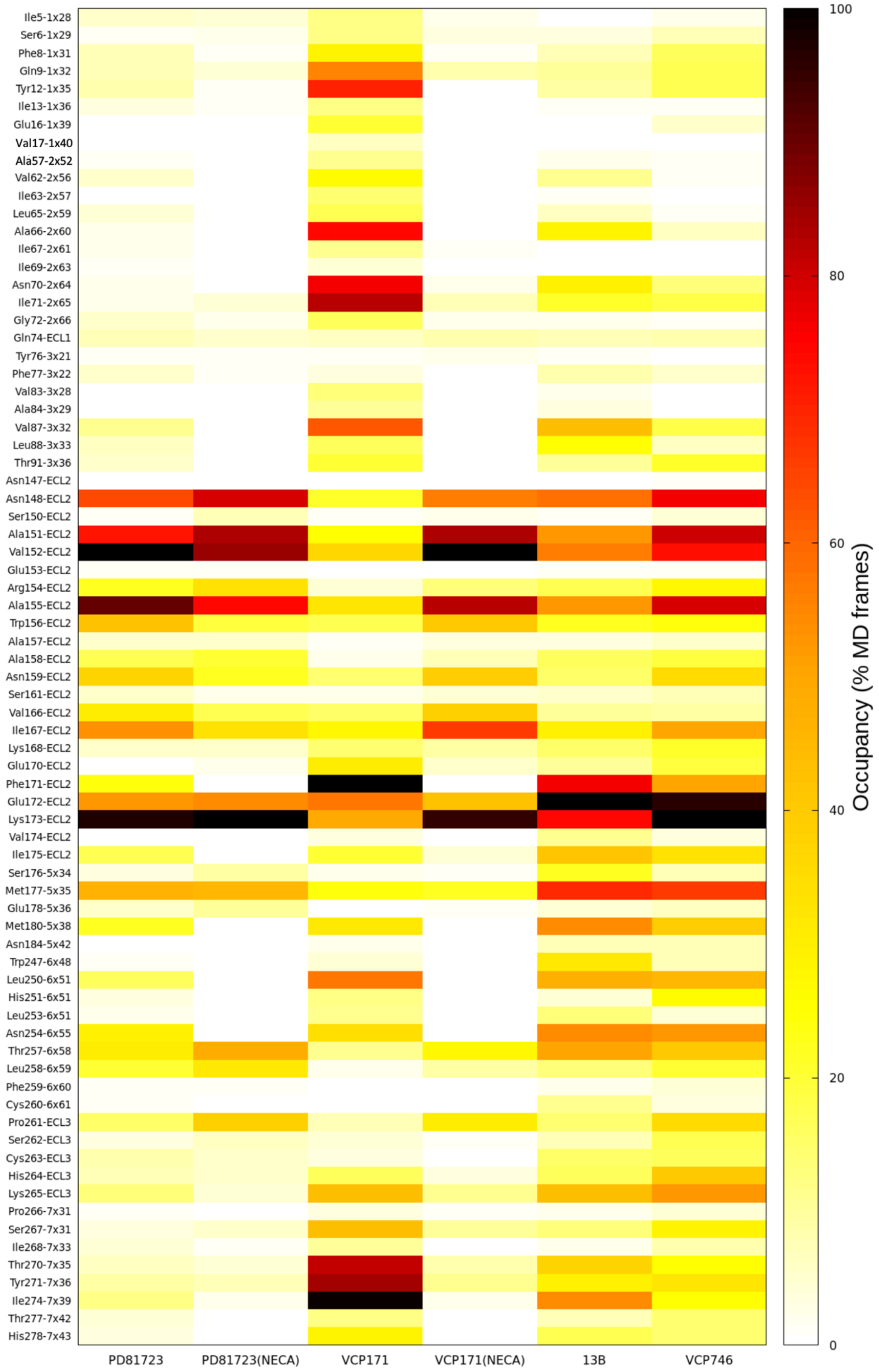


**Figure S1. Ligands - A$_1$R contacts heatmap.** For each A$_1$R residue in each system, the contact occupancy (% MD frames in which the interaction occurred) is normalized for the highest occupancy value.

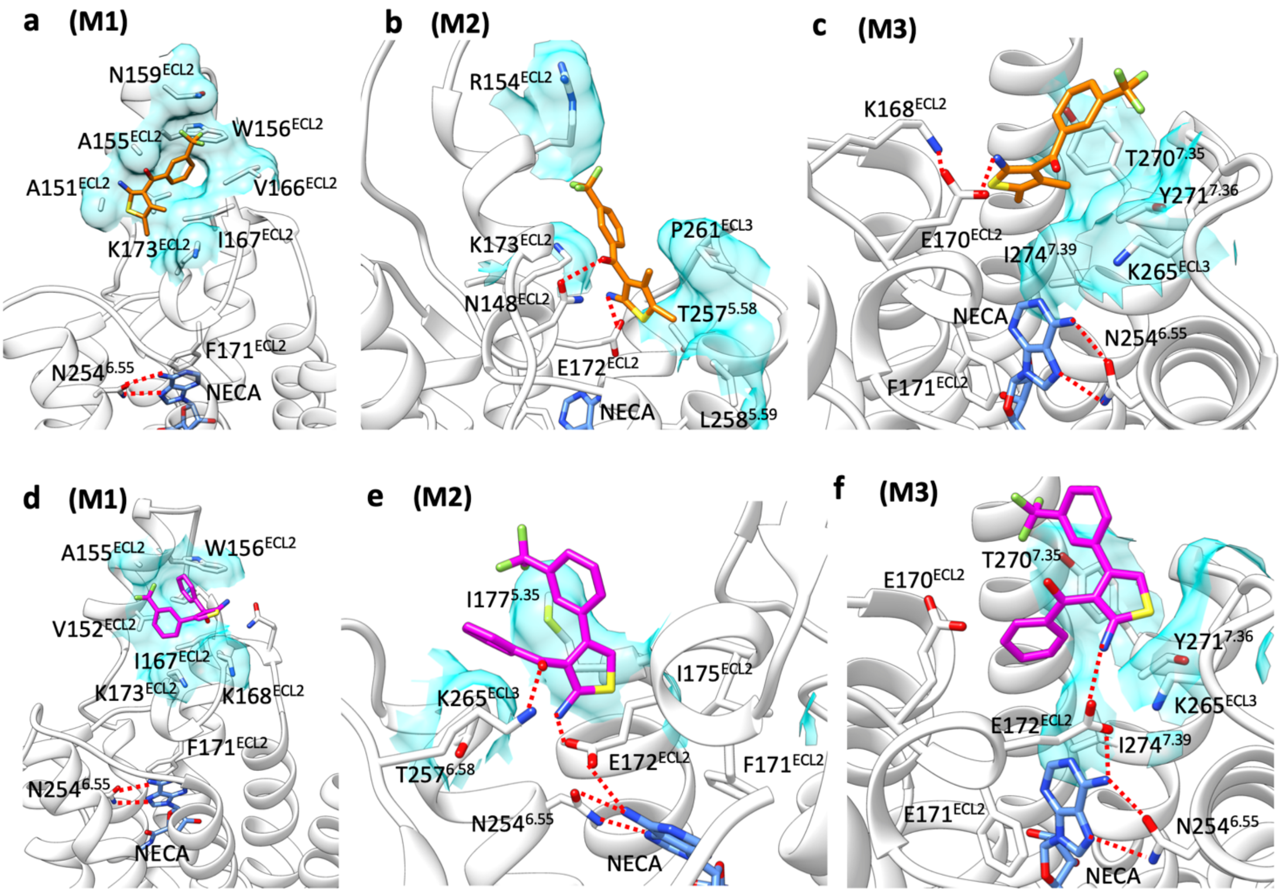


**Figure S2. Representative complexes (microstates) from PD81723 and VCP171 SuMD binding simulations of the occupied $A_1$R. a) - c)** representative configurations from PD81723 (orange stick) metastable macrostates M1-M3 in presence of NECA (blue stick) in the orthosteric site; **d) - f)** representative configurations from VCP171 (magenta stick) metastable macrostates M1-M3 in presence of NECA (blue stick) in the orthosteric site. Hydrophobic contacts are shown as transparent cyan surfaces, hydrogen bonds are reported as dashed red lines.

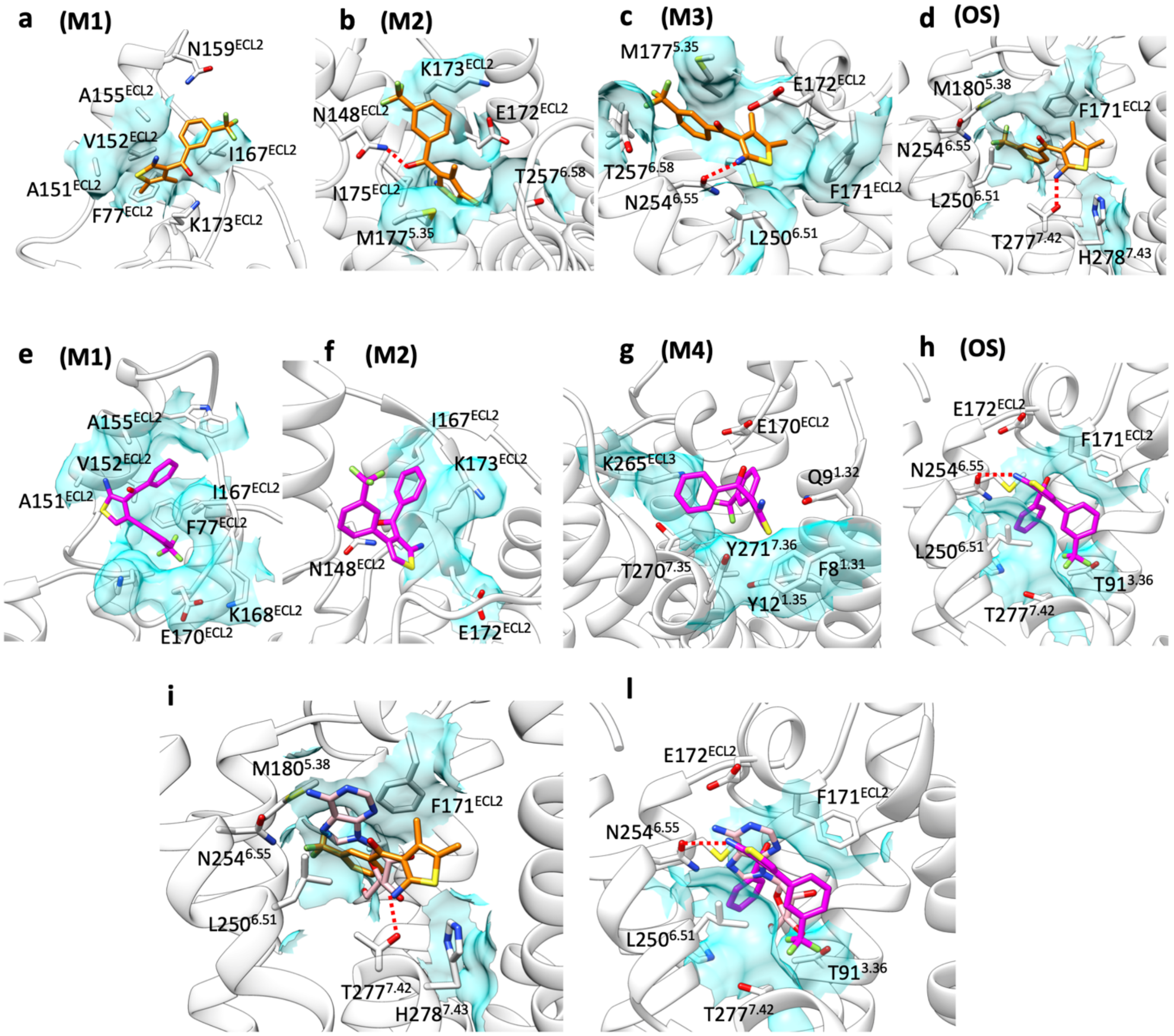


**Figure S3. Representative complexes (microstates) from PD81723 and VCP171 SuMD binding simulations on the unoccupied $A_1R$. a) - d)** representative configurations from PD81723 (orange stick) metastable macrostates M1-M3 and the bound state OS; **d) - h)** representative configurations from VCP171 (magenta stick) metastable macrostates M1, M3, M4 and the bound macrostate OS. **i)** superposition between PD81723 (OS) and adenosine (cryo-EM structure, pink); **l)** superposition between VCP171 (OS) and adenosine (cryo-EM structure, pink). Hydrophobic contacts are shown as transparent cyan surfaces, hydrogen bonds are reported as dashed red lines; in h) TM6 residues are not shown.

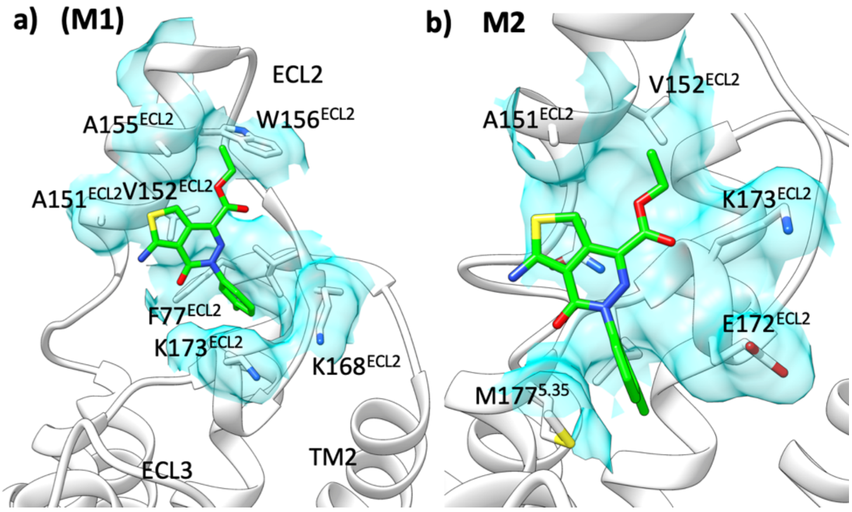


**Figure S4. Representative complexes (microstates) from compound 13B SuMD binding simulations on the unoccupied inactive $A_1R$.** a) Microstate from PD81723 (green stick) metastable macrostates M1; b) Microstate from PD81723 (green stick) metastable macrostates M2. Hydrophobic contacts are shown as transparent cyan surfaces.

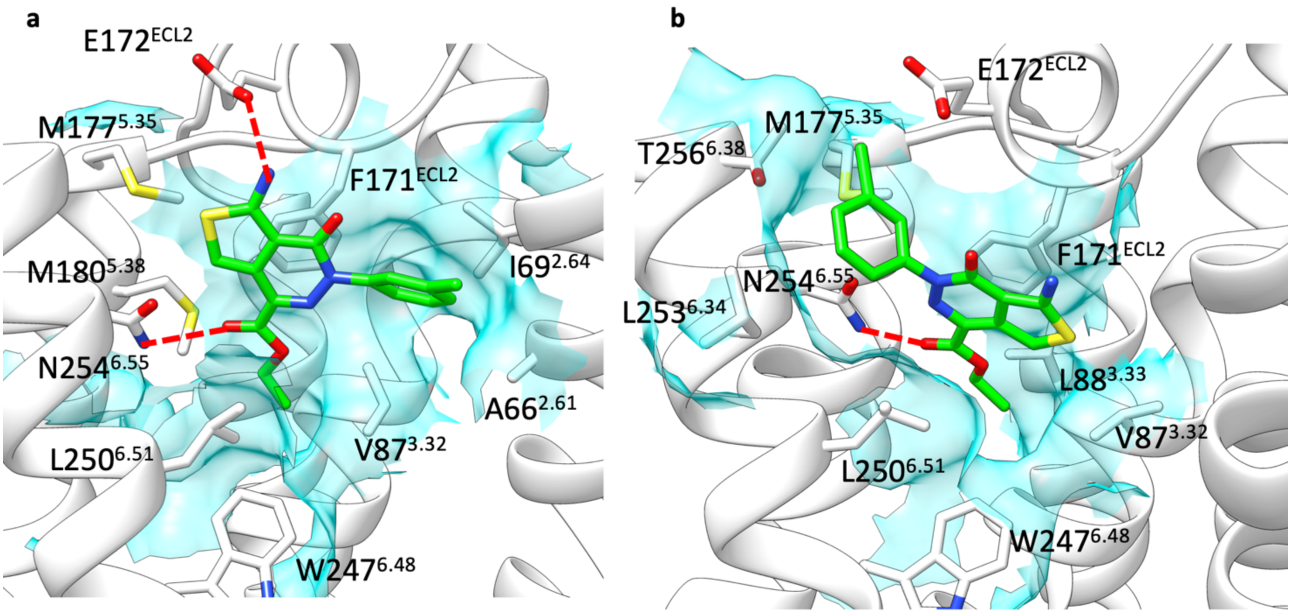


**Figure S5. Representative complexes (microstates) from compound 13B SuMD binding simulations on the unoccupied inactive $A_1R$.** a) and b) Microstates from PD81723 (green stick) orthosteric macrostate OS. Hydrophobic contacts are shown as transparent cyan surfaces, hydrogen bonds are reported as dashed red lines.

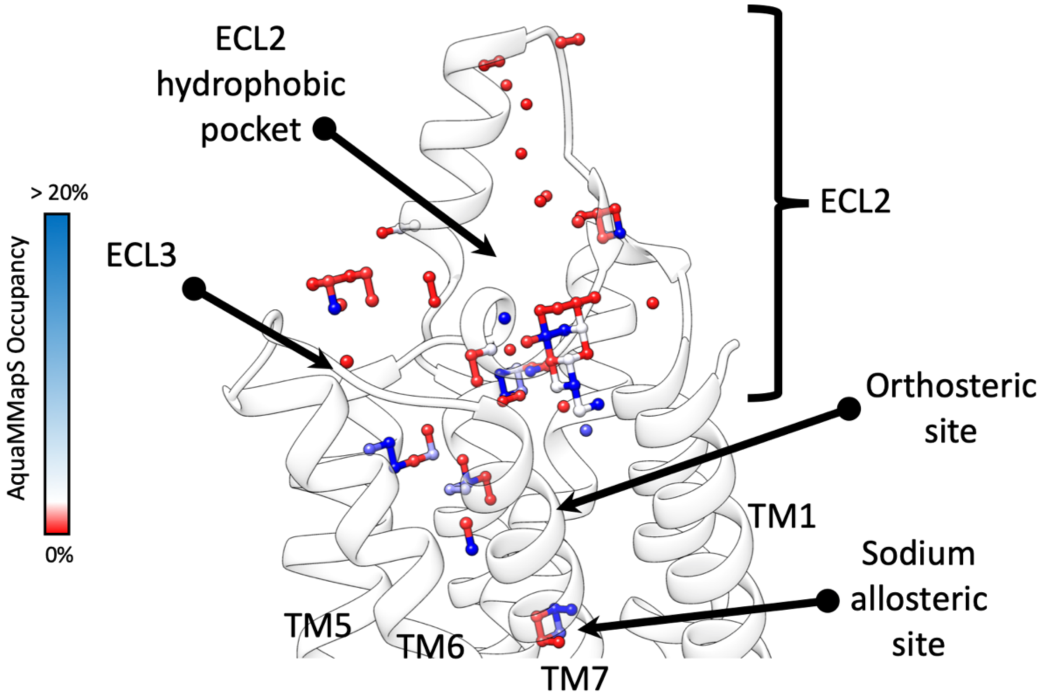


**Figure S6.** Hydrated spots on the $A_1R$ surface (within 4 Å of backbone atoms) according to AquaMMaPS analysis. Beside the allosteric site for the sodium, further region characterized by stabilized water molecules occurred in the orthosteric site and at the ECL2 and ECL3. The hydrophobic pocket within ECL2 (responsible for stable states M1 in Figure 2, Figure 3, Figure S2, Figure S3) shows low tendency to stable hydration.

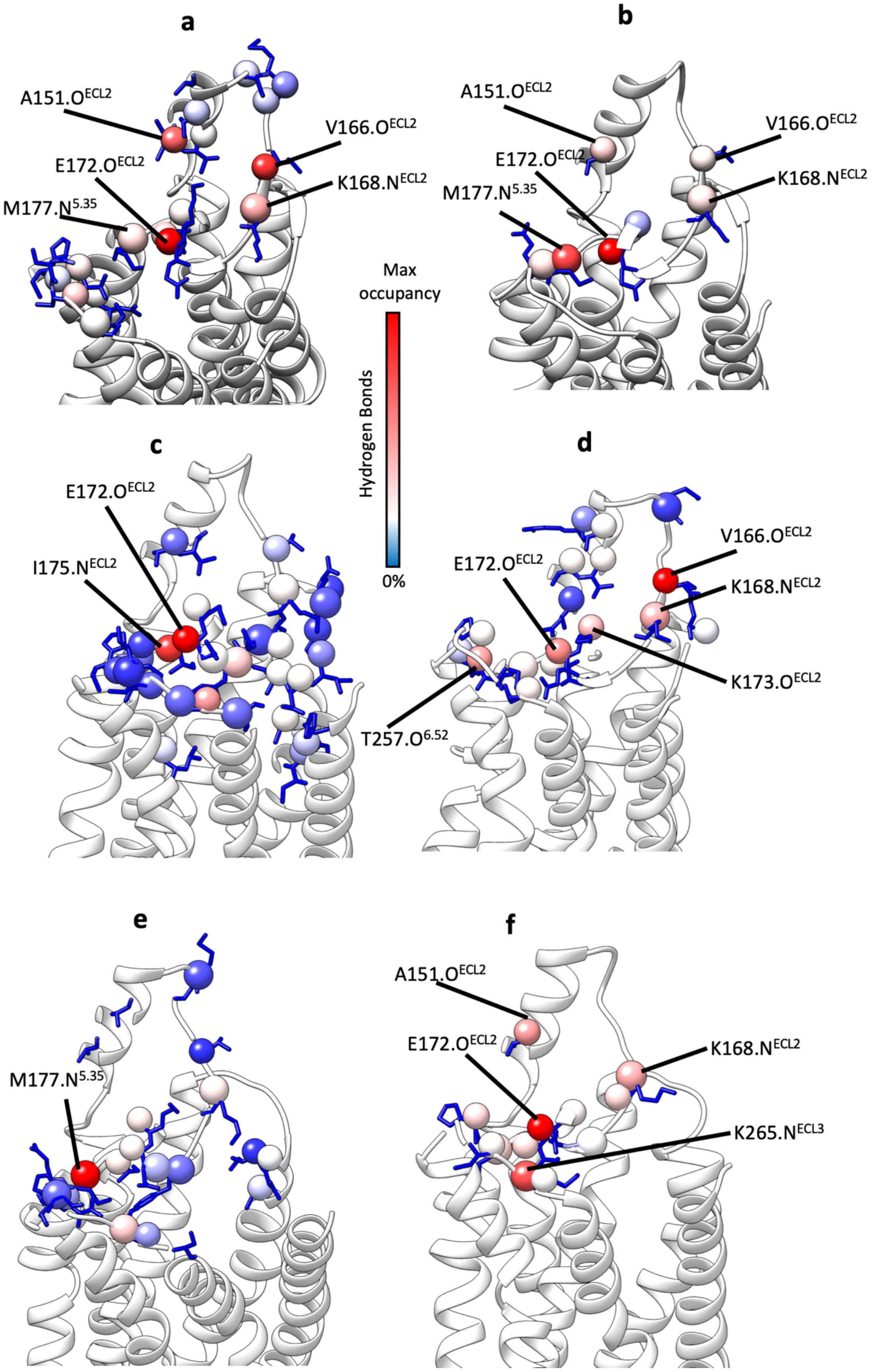


**Figure S7. Main** $A_1R$ backbone atoms involved in hydrogen bonds during simulations. **a**) PD81723-occupied $A_1R$ (NECA bound to the orthosteric site); **b**) PD81723-unoccupied $A_1R$; **c**) VCP171-

occupied $A_1R$ (NECA bound to the orthosteric site); **d**) VCP171-unoccupied $A_1R$; **e**) compound 13B-unoccupied inactive $A_1R$; f) **b**) CP746-unoccupied $A_1R$.

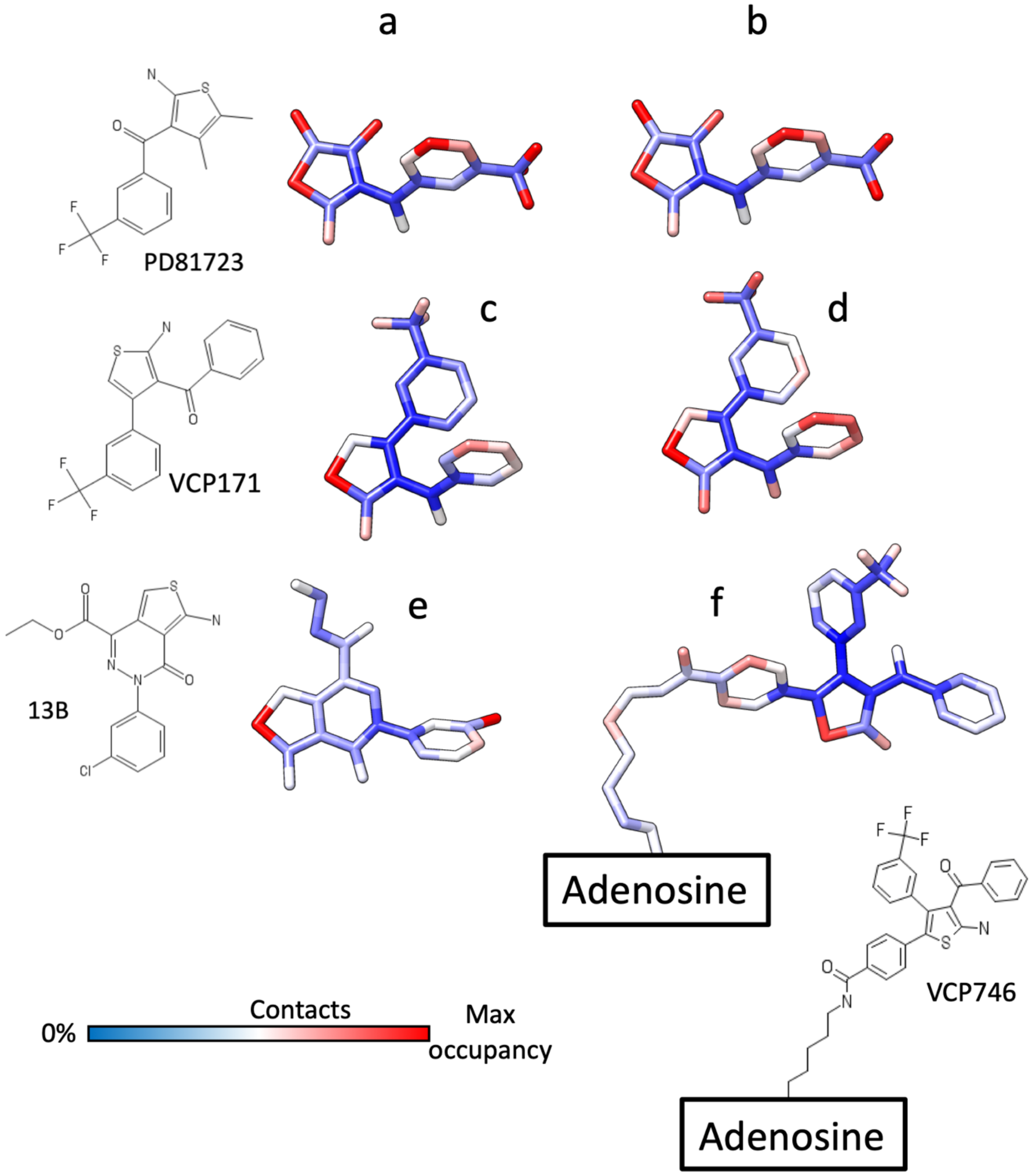


**Figure S8. Ligands' atoms involved in contacts with $A_1R$ during SuMD simulations. a**) PD81723-occupied $A_1R$; **b**) PD81723-unoccupied $A_1R$; **c**) VCP171-occupied $A_1R$; **d**) VCP171-occupied $A_1R$; **e**) Compound 13B-unoccupied inactive $A_1R$; **f**) VCP746-unoccupied $A_1R$. Atoms are colored according to the total percentage of MD frames (occupancy) in which they were involved in contacts with the receptor. Among the simulated ligands, compound 13B was the less involved at the level of the 2-aminogroup.

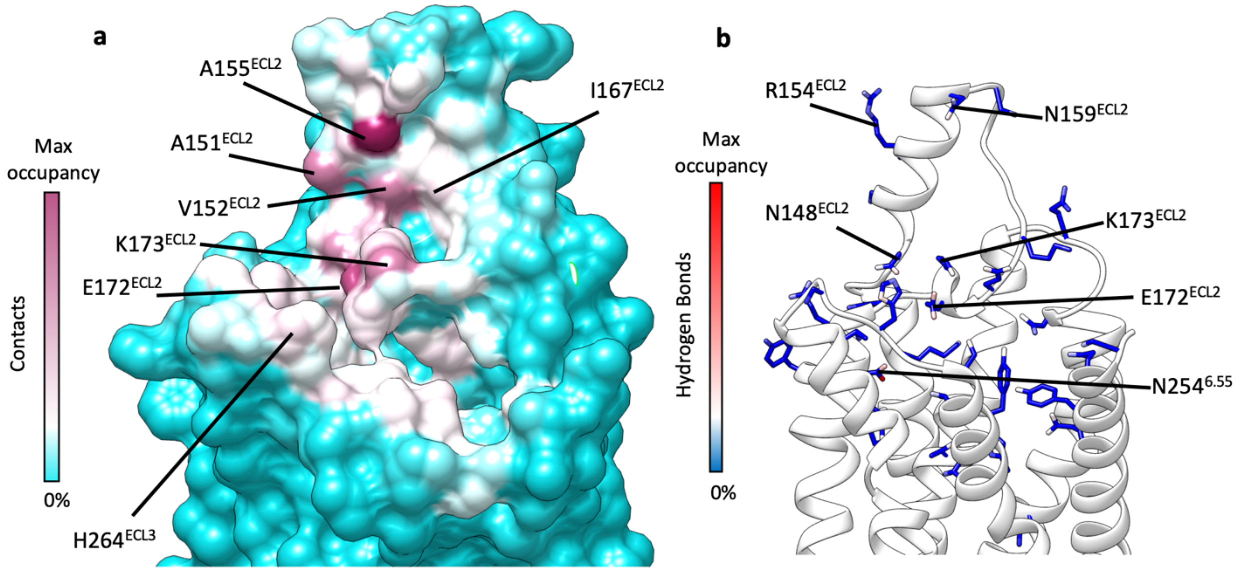


**Figure S9. a)** $A_1$R-VCP746 contacts plotted on the receptor surface and colored according to the occupancy (% MD frames in which a contact was present); **b**) $A_1$R side chain atoms that formed hydrogen bonds with VCP746 and colored according to the occupancy (%MD frames in which a hydrogen bond was present).
